# Bioimpedance-assisted characterization of cardiac electroporation and anisotropic homogenization by pulsed field ablation

**DOI:** 10.64898/2026.03.18.712769

**Authors:** Edward J. Jacobs, Pedro P. Santos, Sam S. Parizi, Simon N. Dunham, Rafael V. Davalos

## Abstract

**Objective:** Pulsed field ablation (PFA) relies on irreversible electroporation to create nonthermal cardiac lesions, yet real-time indicators of electroporation progression and validated lethal electric field thresholds remain limited. This study aimed to develop a bioimpedance-based metric for real-time monitoring of cardiac electroporation, evaluate the impact of myocardial anisotropy under electroporation conditions, and derive waveform-specific lethal electric field thresholds.

**Introduction:** Current PFA procedures lack direct intraoperative feedback on lesion formation, and uncertainty remains regarding the role of myocardial fiber orientation in shaping electric field distributions. Because electroporation dynamically alters tissue electrical properties, monitoring these changes during treatment may improve prediction of ablation outcomes.

**Methods:** PFA was delivered to fresh *ex vivo* porcine ventricular tissue using clinically relevant and energy-matched waveforms with pulse widths from 1 to 100 µs. Inter-burst broadband electrical impedance spectroscopy was performed using a low-voltage diagnostic waveform to quantify burst-resolved impedance changes. Lesions were visualized using metabolic staining, then finite element models incorporating nonlinear electroporation-dependent conductivity were used to compare anisotropic and homogenized electric field distributions. Lethal electric field thresholds were estimated by fitting simulated contours to measured lesion areas and validated using uniform electric fields generated by a parallel electrode array.

**Results:** Across all waveforms, impedance measurements showed a rapid initial decrease followed by stabilization, indicating early electroporation saturation. Burst-to-burst percent change in impedance slope provided a consistent, waveform-agnostic metric of electroporation progression. Lesion morphology was not systematically influenced by fiber orientation, and modeling demonstrated that electroporation-induced conductivity increases homogenized tissue anisotropy. Lethal electric field thresholds increased with decreasing pulse width, ranging from 517 ± 46 V/cm (100 µs) to 1405 ± 55 V/cm (1 µs), and were validated under uniform field conditions.

**Conclusion:** Bioimpedance-assisted monitoring enables real-time assessment of cardiac electroporation, while electroporation-induced homogenization supports simplified modeling and standardized PFA treatment design.

## 1. Introduction

Catheter-based pulsed field ablation (PFA) has transformed the treatment of atrial fibrillation, utilizing high-amplitude, low-energy pulsed electric fields (PEFs) to ablate or isolate asynchronous cardiac tissue^1–3^. PEFs generate an ionic current within the tissue that induces an electric potential across the plasma membrane of cells, as charges of opposite polarity accumulate on each side of the membrane. Once the induced transmembrane potential reaches a critical voltage (∼0.258 V)^4^, the membrane permeabilizes through the formation of nanoscale defects (electroporation)^5^. If persistent, the cells within a critical electric field (EF) threshold die through loss of homeostasis, termed irreversible electroporation (IRE)^6,7^. The selective, nonthermal mechanism of IRE minimizes injury to peri-atrial structures such as the esophagus and phrenic nerve and reduces the risk of pulmonary vein stenosis compared to radiofrequency or cryothermal ablation^8^. Recent multicenter trials have confirmed that PFA achieves comparable or superior acute efficacy relative to thermal ablation while significantly reducing complications^9–16^. However, despite the clear procedural and safety advantages of PFA, its ongoing clinical translation still faces important technical challenges.

There is currently no clinical tool to evaluate the completeness of lesions. The absence of real-time, intraoperative monitoring of electroporation progression limits feedback on lesion completeness during treatment, leading to potential gaps in transmural coverage or unnecessary repeat applications. While methods are being developed to visualize the ablative footprint, field tags are purely geometric and do monitor electroporation generation itself^17^, which may be different between patients and in diseased tissues. This means that verifying acute treatment success still requires post-treatment electrical mapping using expensive mapping systems that are not common outside of the United States, with follow-up surveillance to validate chronic disease control. Further, acute post-treatment electrical mapping may not detect cardiac tissue temporarily “stunned” by reversible electroporation^18^, leading to undertreatment and eventual pulmonary vein reconnection. If higher treatment energies are applied to compensate for potential stunning, over-treatment may increase complications through excessive bubbles, hemolysis, electrolytic effects, or thermal damage to sensitive structures. Moreover, the diversity of commercially implemented waveforms and catheter geometries has produced substantial variability in lesion depth and continuity, complicating the establishment of universal treatment parameters^19,20^. Only a few commercial catheters measure impedance ^21–23^, but typically to gauge probe conformation or tissue contact, not the extent of electroporation.

The impact of tissue composition and geometry with electroporation effects is also still being elucidated. The depth and uniformity of the PFA lesion are governed by both the local electric field distribution and the waveform-specific electric field threshold for electroporation. Computational modeling is essential during therapy development for visualization of expected field distributions, ensuring that the entire myocardial wall is exposed to lethal field strengths while sparing adjacent critical structures. Previous studies have largely inferred anisotropic effects from computational models, suggesting that fiber orientation might influence the electric field distribution during PEF application after forcing the electric field to perfectly align with fibers spanning the entire domain^24^. However, experimental measurements only observe differences when the electric field is precisely aligned with fiber orientation and disappear when deviating by more than 10-15 degrees^25,26^. While these results are supported in isotropic skeletal muscle fibers^27,28^, cardiac tissue is not as unidirectional as skeletal muscle and does not have large areas of parallel fiber orientation^29^, especially in atrium and diseased tissue. Further, there is limited experimental evidence of anisotropy under electroporation conditions. Nevertheless, as the cell membranes become permeabilized through electroporation, local tissue electrical conductivity increases nonlinearly, significantly altering the electric field geometry^30^. Recent work suggests that the dynamic changes in tissue electrical properties during electroporation may homogenize tissue impedance, obviating any potential directional dependence^30,31^. Achieving consistent and durable lesions, therefore, requires a quantitative understanding of how waveform construction interacts with anisotropic cardiac tissue to produce regions of electroporation.

Here, we implemented inter-burst broadband electrical impedance spectroscopy (EIS) during PFA treatments to elucidate the evolution of cardiac electroporation across multiple clinical waveforms. We then proposed methods for utilizing tandem bioimpedance as a metric for evaluating treatment saturation. Following, we demonstrate that electroporation homogenizes tissue bioimpedance to mask potential cardiac directional dependency, using measured IRE areas with characterized fiber orientations and non-linear anisotropic electroporation modeling. These results advance our understanding of electroporation evolution during cardiac PFA, provide validated, transparent cardiac lethal electric field thresholds, and interrogate assumptions underlying cardiac PFA modeling. Together, these data enable better predictive modeling, standardized waveform selection, and more reproducible clinical outcomes across patient populations and device platforms.

## 2. Results

### 2.1. Experimental setup and determination of treatment parameters using a mimicking computational model

To determine pulse parameters for generating measurable IRE lesions in cardiac tissue, we implemented an electroporation- and thermal-dependent computational model of our experimental setup. As the applied voltage would affect the electric field that influences the rate of electroporation within the tissue, we aimed to determine a spacing-normalized voltage that would produce measurable areas for each of the waveforms previously characterized^30^: 1-1-1-1-μs x 50, 5-5-5-5-μs x 10, 10-10-10-10-μs x 10, and 100-μs. Two sharp-tip 1-cm exposure monopolar electrodes were spaced 1 cm apart and inserted 2 cm deep within a geometry replicating a sliced porcine ventricle (**Figure 1A**). The non-linear increase in electrical tissue conductivity due to electroporation was defined as a function of local electric field intensity based on previously characterized curves within 37°C porcine ventricular tissue for these waveforms (**Figure 1B**)^30^. Further, to estimate the expected lesions within this experimental set up, we also defined the lethal electric field thresholds from previously characterized human cardiomyocytes within an *in vitro* tissue mimic for these waveforms (**Figure 1C**) ^30^. We then iteratively increased the voltage and simulated the expected region of IRE. The simulations suggest that 2500 V would produce similar electric field distributions, with slight variability due to differences between the non-linear conductivity curves (**Figure 1D)**. Further, the 2500 V application should also produce measurable lesions for each of the proposed waveforms, with differences due to the expected lethal electric field thresholds (**Figure 1E**). These results provide a singular experimental voltage to produce measurable lesions across the range of PFA waveforms tested, while maintaining relatively equal electric field distributions and thermal heating.

**Figure 1.**
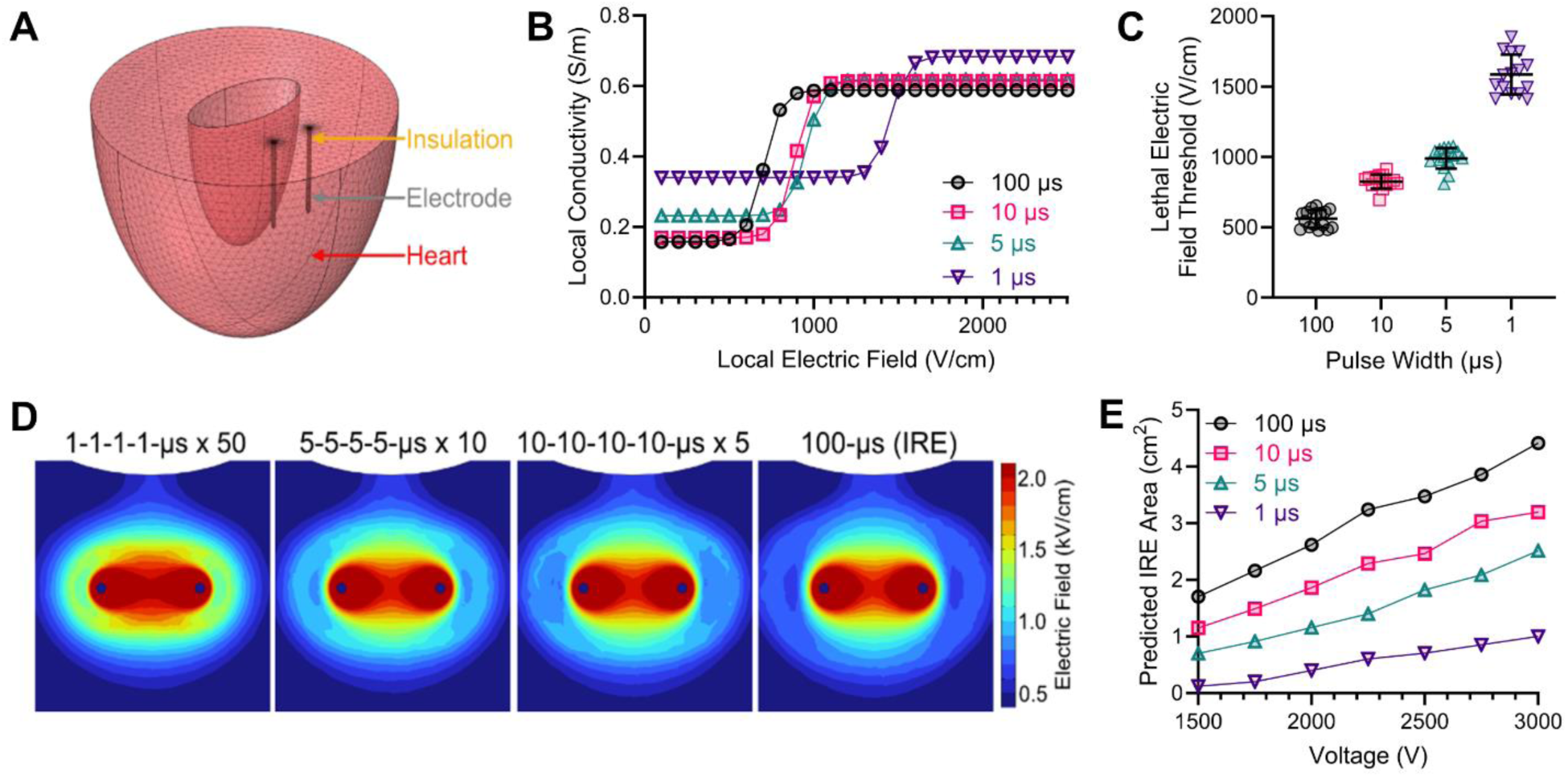
Electroporation finite element model of the cardiac experimental setup. **A)** Two sharp-tip 1-cm exposure monopolar electrodes spaced 1 cm apart and inserted 2 cm deep within a half porcine ventricle (COMSOL^TM^ v6.4). **B)** Defined cardiac electroporation-dependent electrical conductivity and **C)** previously characterized IRE thresholds for *in vitro* human cardiac cells for each of the experimental waveforms^30^; mean ± SD; n=16. **D)** Electric field distributions on a cut plane through the middle of the electrodes for each of the waveforms with 2500 V between 1-cm center-to-center electrode spacing. **E)** Predicted irreversible electroporation (IRE) lesion area from different applied voltages, using the finite element models with average waveform-specific conductivity curves and lethal thresholds.

### 2.2. Inter-burst bioimpedance monitoring reveals rapid cardiac electroporation saturation during pulsed field ablation across multiple waveforms

After selecting the treatment voltage, we then delivered PFA treatments within fresh *ex vivo* porcine ventricular tissue using the same electrode configuration from modeling^30,32^. To gauge the level of disruption, we measured the electrical impedance spectra using our novel inter-burst EIS technique, termed Fourier Analysis Spectroscopy (FAST)^33,34^. The diagnostic FAST waveform is a 12 V custom bipolar chirp consisting of both high-frequency (2-48-2-48-µs x 50 cycles) and low-frequency (250-1-250-499-µs x 5 cycles) concatenated signals used to perform EIS over a wide frequency range (1kHz – 0.5MHz) (**Figure 2A**). The cell membrane acts as an insulator, and at low frequencies and low voltages, electric currents cannot easily pass through it and instead flow around the cells in the extracellular fluid, resulting in a higher electrical impedance (**Figure 2B**). However, as the frequency of the waveform increases, the current can pass through the cell membrane and the highly conductive cytoplasm inside the cells, lowering the overall tissue impedance. The shift in how electric current interacts with the cellular structure as the frequency increases, from being blocked by the membrane to passing through it, causes a characteristic change in the tissue’s electrical properties and the Maxwell-Wagner effect (i.e., beta dispersion)^35,36^. The formation of nanopores in cell membranes during electroporation delivery facilitates intra- and extra-cellular ion exchange, effectively reducing the ability of the cell membrane to act as an insulator. The FAST diagnostic waveform is delivered at a low voltage (12 V) to avoid inducing further impedance changes while sensing the non-linear electroporation effects^33^. FAST diagnostic measurements were performed directly before treatment and then between each burst to evaluate the evolution of electroporation effects during PFA treatments. Over the course of pulse delivery, the low-frequency impedance drops, and when sufficient pulses have been delivered to fully electroporate the tissue, the low-frequency impedance resembles the high-frequency impedance, at which point additional permeabilization will not occur^33,37^. Every cardiac tissue treatment across each waveform displayed the disruption of the cardiomyocyte cellular membrane, with a decrease in low- and high-frequency impedance over treatment (**Figure 2C**).

**Figure 2.**
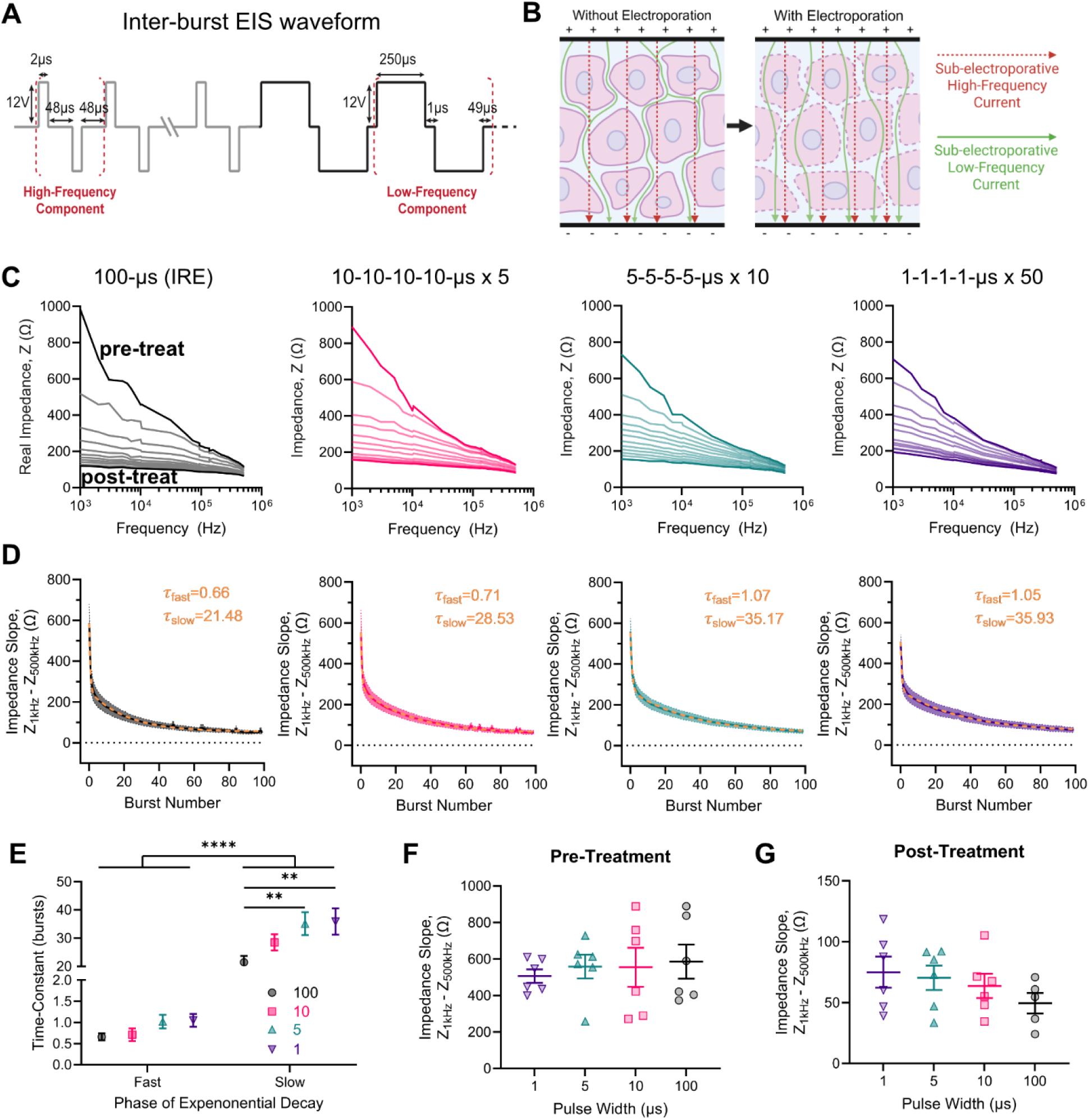
Inter-burst electrical impedance spectroscopy (EIS) during pulsed field ablation delivery. **A)** Inter-burst EIS waveform consisting of low-frequency and high-frequency components for broadband 0.1 to 500 kHz EIS. **B)** Before electroporation, low-voltage low-frequency displacing current cannot pass through the cell membrane, while higher frequency conduction currents more easily pass. Electroporation disrupts the cell membrane impedance, allowing low-voltage low-frequency displacement currents to pass through the membrane. **C)** Representative pre-treatment EIS, inter-burst EIS (every 10 bursts plotted), and post-treatment EIS measurements for each of the waveforms over 100 bursts at 1 Hz. **D)** Difference in low-frequency and high-frequency impedance (slope) for pre-treatment (0), between applied bursts (1-99), and post-treatment (100) for each waveform; mean ± SEM; n = 6; data fit with an unrestricted 2-phase exponential decay. **E)** Time constants for the 2-phase exponential decay; mean ± SEM; n = 6; Two-way ANOVA with multiple comparisons within each row and within each column (** *p <* 0.01, **** p < 0.0001). **F)** Pre-treatment and **G)** post-treatment impedance slope for each waveform; One-way ANOVAs with Tukey’s post hoc (no significance); mean ± SEM; n = 6.

As the difference in low- and high-frequency impedance can be attributed to the cardiac cell’s ability to act as an electrical insulator, we quantified the evolution of this difference (“slope”) for every burst (**Figure 2D**). For each waveform, we observed a large initial drop in bioimpedance slope, followed by a more gradual decrease for the rest of treatment. Therefore, we fit an unconstrained 2-phase exponential decay to account for the initial electroporation of the intact membrane and the continued disruption of already permeabilized cells. The 2-phase exponential decay was unconstrained to allow for the best fit of both exponential curves and the transition point between both phases. All curves had goodness-of-fit R^2^ values above 0.6500 (p < 0.0001). The 1^st^ phase fast time-constants were determined for IRE (0.66 ± 0.083 bursts), 10-μs (0.71 ± 0.15 bursts), 5-μs (1.02 ± 0.16 bursts), and 1-μs (1.05 ± 0.15 bursts). The 2^nd^ phase was also determined for IRE (21.48 ± 2.21 bursts), 10-μs (28.53 ± 2.88 bursts), 5-μs (35.17 ± 4.05 bursts), and 1-μs (35.93 ± 4.65 bursts), and each waveform had a significantly longer 2^nd^ phase time constant than 1^st^ phase time constant (**Figure 2E**, all p<0.0001). As the pre-treatment non-electroporated slopes were not significantly different across pulse width groups (**Figure 2F**), we do not attribute these trends to innate differences in the cardiac tissue samples. Further, there were no significant differences in post-treatment impedance slope between the groups, but there was a negative trend with increasing pulse width (**Figure 2G**). Concomitantly, the transition point between the two phases was determined for IRE (1.45 ± 0.58 bursts), 10-μs (1.40 ± 0.37 bursts), 5-μs (1.08 ± 0.08), and 1-μs (1.45 ± 0.30). Together, the rapid decrease in tissue impedance across burst 1, followed by the transition to a slow exponential decay between bursts 1 and 2, supports that the tissue is initial permeabilized, then subsequent pulses maintain permeabilization while further disrupting the membrane impedance to a lesser extent.

### 2.3. Proposed method for inter-burst monitoring of cardiac PFA saturation

While analysis of EIS post-treatment can interrogate how the tissue reacts to electroporative electric fields, it does not provide real-time feedback for informing treatment completion. For real-time processing of bioimpedance data, metrics are needed to gauge the extent of permeabilization. We propose using percent change in burst-to-burst in impedance slope as a gauge of the rate of membrane permeabilization. For each burst (bi), the percent change in impedance slope relative to the immediately preceding burst (bi-1) was calculated and tracked over the course of treatment for each waveform (**Figure 3A**). Following burst 1, the percent change in impedance slope decreased monotonically with additional pulse delivery and approached near-zero values within the early portion of treatment for all pulse widths. This trend was consistent across waveforms despite differences in absolute impedance values and pulse structure, indicating a shared temporal profile of diminishing impedance evolution. Notably, no waveform exhibited secondary divergence in burst-to-burst percent change once the initial decrease occurred, with the observed oscillations due to small shifts in measured nominal impedance as seen in Figure 2D.

**Figure 3.**
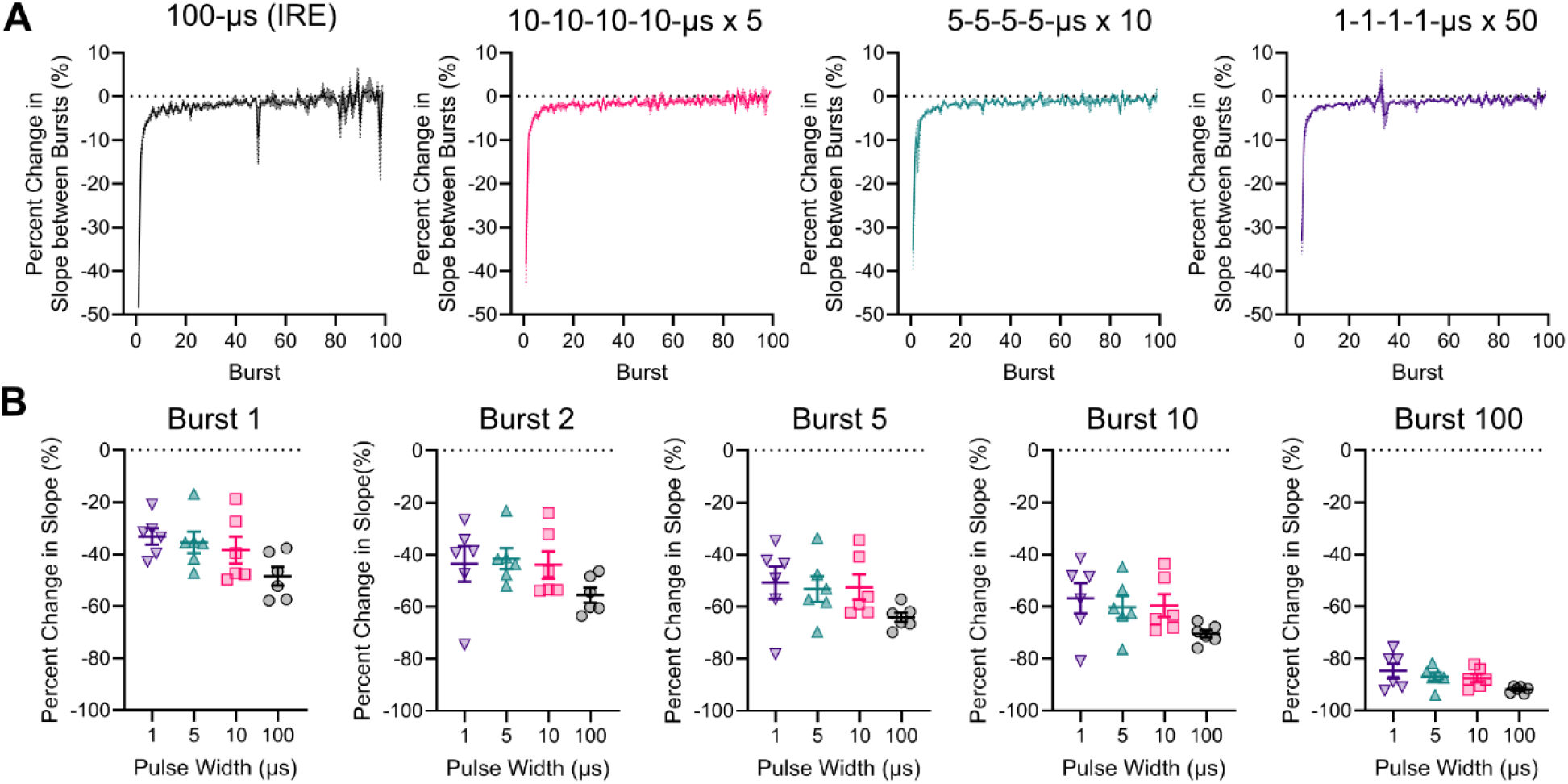
Method for inter-burst quantification of PFA saturation. **A)** Percent change in impedance slope across each burst (b_i_) for each waveform; mean ± SEM; n = 6**. B)** Percent change in impedance slope from pre-treatment at different bursts (b_i_ – b_0_); mean ± SEM; One-way ANOVAs with Tukey’s post hoc (no significance); n = 6.

To evaluate cumulative electroporation effects, the percent change in impedance slope relative to pre-treatment was also computed for each burst (**Figure 3B**). While early bursts produced rapid divergence from baseline, subsequent bursts resulted in progressively smaller incremental changes. No significant differences were observed between waveforms at matched burst numbers, indicating that the relative progression toward impedance stabilization followed similar trajectories across pulse widths. Together, these results suggest that percent changes in impedance slope provide a consistent, waveform-agnostic measure of electroporation progression, with early bursts dominating the measurable impedance evolution and later bursts producing minimal additional change.

### 2.4. Homogenization of the local conductivity removes the impact of anisotropy on pulsed field ablation

The measured decrease in impedance slope suggests the attenuation of membrane capacitance, which directly contributes to electrical anisotropy in tissue^35^. To assess whether myocardial fiber orientation influences PFA outcomes under electroporation conditions, we analyzed lesion formation in cardiac tissue regions with distinct fiber alignments relative to the applied electric field. Following treatment, tissue sections were stained with 2,3,5-tetrazolium chloride (TTC) to identify metabolically inactive regions corresponding to IRE (**Figure 4A**)^20,32^. Fiber orientation was independently characterized in a blinded manner and categorized as predominantly aligned with the applied field (−30° to 30°) or predominantly perpendicular (60° to 120°), with only 10 of lesions presenting clear fiber orientation within these two categories. Despite differences in underlying fiber orientation, TTC staining revealed contiguous lesions with comparable shapes and extents across orientations and waveforms. No systematic elongation or directional bias in lesion morphology was observed between aligned and perpendicular regions. The formation of IRE lesions also supports the voltage chosen from initial computational modeling and the indirect monitoring of electroporation progression from FAST bioimpedance measurements.

**Figure 4.**
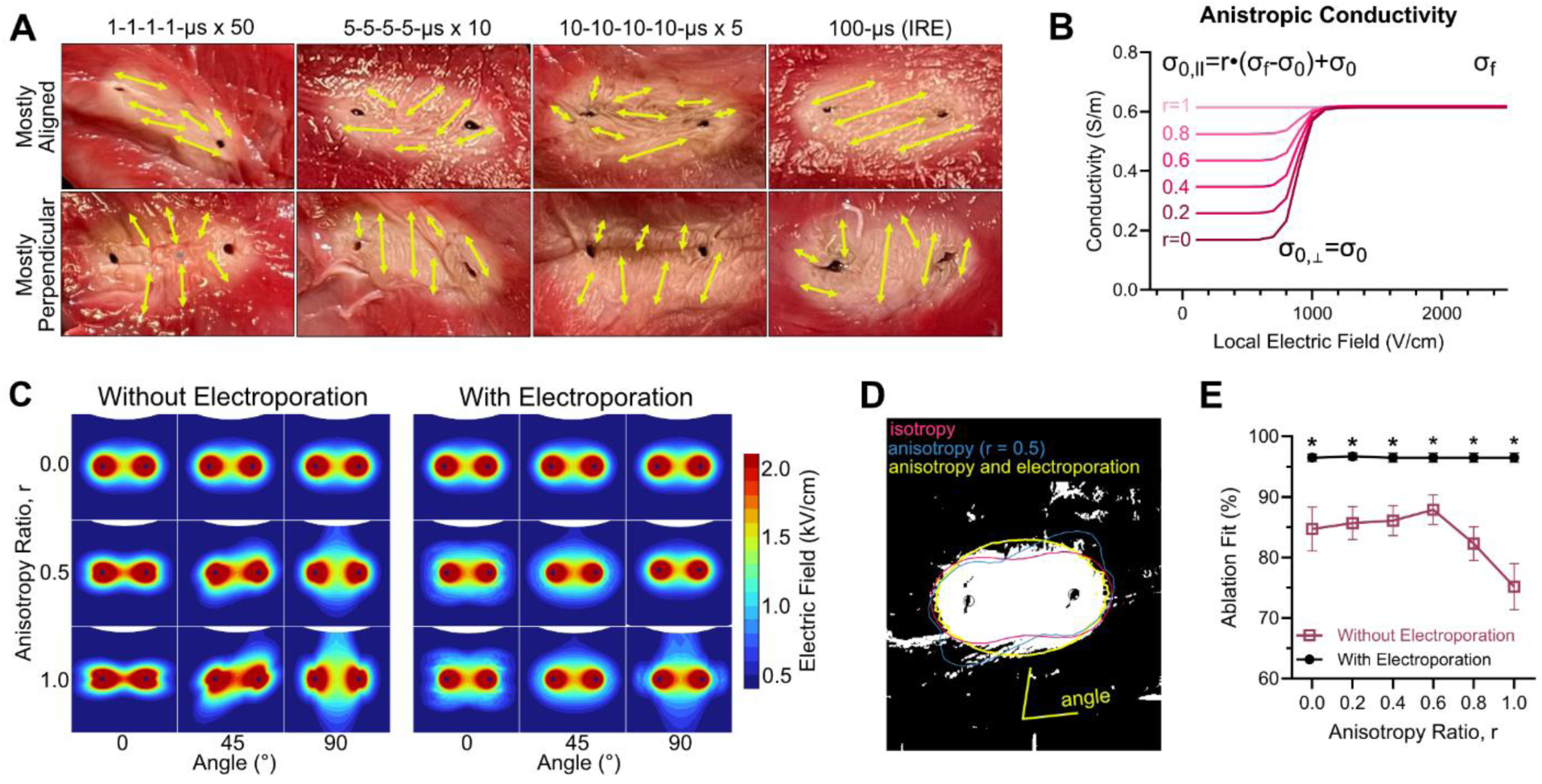
Analysis of lethal electric field contour similarity with isotropy and electroporation effects. **A)** Metabolic 2,3,5-Triphenyl tetrazolium chloride (TTC) staining of cardiac tissue slices ∼ 4 hours after pulsed field ablation (red indicates metabolically active tissue). Fiber orientation drawn after unbiased blind characterization with mostly aligned -30-30° and mostly perpendicular 60-120°. **B)** Conductivity curves for the parallel and perpendicular directions to create anisotropic tissue. **C)** Finite element simulations (COMSOL^TM^) of the electric field distribution in the cardiac tissue under electroporation with different anisotropic ratios and fiber orientations. **D)** Representative 8-bit image conversion with auto-color-thresholding (ImageJ Fiji) and overlayed electric field contours (COMSOL^TM^) for isotropic, medium anisotropy (r=0.5), and anisotropy (r=0.5) with electroporation. **E)** Lesion and area-matched contour fits for different anisotropic ratios with and without electroporation areas; mean ± SEM; Student T-tests at each anisotropic ratio (* p < 0.0001); n = 10.

These experimentally measured lesion areas were subsequently used to evaluate electric field distribution fits generated under anisotropic and homogenized conductivity assumptions. To model anisotropic tissue behavior, direction-dependent conductivity curves were assigned parallel and perpendicular to the local fiber orientation (**Figure 4B**). These curves incorporated experimentally measured nonlinear conductivity increases associated with electroporation and an anisotropic ratio (r) to increase the conductivity aligned parallel with the fiber orientation. Electric field distributions were computed using both anisotropic conductivity tensors (**Figure 4C**). Across all waveforms, the predicted lethal contours from the homogenized model closely matched those generated using isotropic conductivities, with minimal spatial deviation.

To quantitatively assess agreement between experimental lesion areas and modeled predictions, lethal electric field thresholds were fit using all anisotropic and homogenized field distributions (**Figure 4D**). For each lesion, the fiber angle was defined within the COMSOL simulation to generate electric field distributions with isotropic conductivity, anisotropic conductivity, and anisotropic conductivity with electroporation. The contour with the same internal area as the lesion was then overlayed with the probe insertion sites to calculate the percentage overlap between the lesion and contour shape. Fits generated using homogenized conductivity fields yielded stable threshold estimates and low residual error across all fiber orientations (**Figure 4E**), with no systematic bias attributable to alignment. Incorporating anisotropy did not significantly improve the quality of the fits or reduce variability in the estimated lethal thresholds. Together, these results demonstrate that electroporation-induced conductivity changes dominate the electric field distribution during pulsed field ablation, resulting in effective homogenization of tissue electrical properties and minimizing the influence of myocardial fiber orientation on lesion formation.

### 2.5. Lethal pulsed field ablation threshold fits using homogenized tissue electric field distributions

Since electroporation homogenizes tissue anisotropy, we could then overlay the simulated electric field contours for all the cardiac lesions where clear fiber orientations were not obtained. Experimentally measured IRE areas were quantified from TTC-stained cardiac tissue slices and overlaid with simulated electric field contours generated using homogenized conductivity curves for each waveform (**Figure 5A–B**). The electric field contour corresponding to the measured lesion boundary was then used to estimate the lethal electric field threshold. Thresholds significantly increased monotonically with decreasing pulse width, with 517 ± 47 V/cm for 100-µs pulses, 655 ± 120 for 10-10-10-10-μs x 5 pulses, 833 ± 84 for 5-5-5-5-μs x 10 pulses, and 1405 ± 42 V/cm for 1-1-1-1-µs x 50 pulses (**Figure 5C–D**; **Table 1**). Fits using homogenized electric field distributions produced consistent threshold estimates with low variability across samples, and no systematic deviations were observed between modeled contours and experimentally measured lesion geometries.

**Figure 5.**
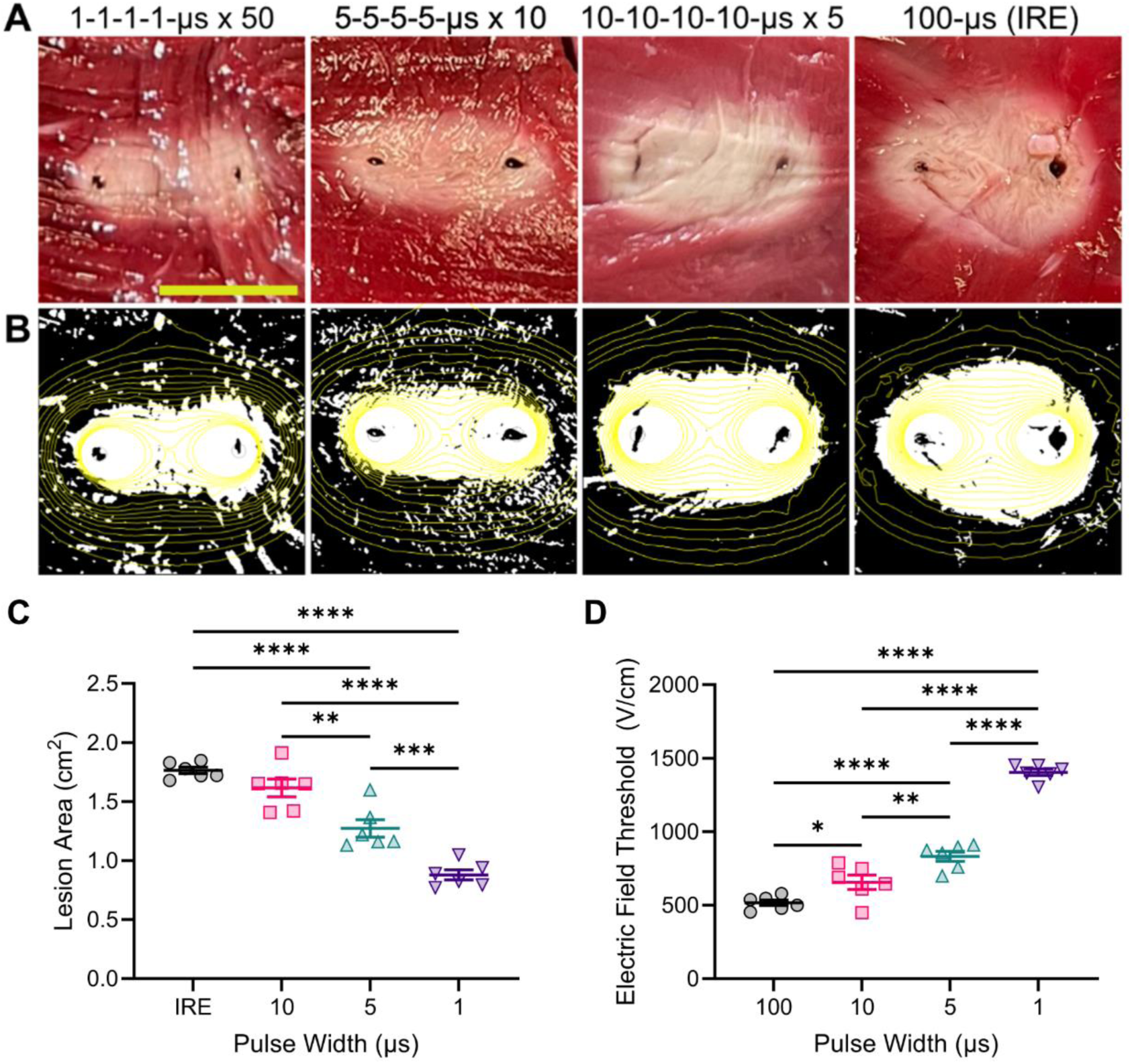
Cardiac pulsed field ablation areas and lethal electric field threshold calculations. **A)** Metabolic 2,3,5-Triphenyl tetrazolium chloride (TTC) staining of cardiac tissue slices ∼ 4 hours after pulsed field ablation (red indicates metabolically active tissue). **B)** 8-bit image conversion with auto-color-thresholding (ImageJ Fiji) and overlayed waveform-specific electric field contours (COMSOL^TM^). **C)** Measured irreversible electroporation lesion areas and **D)** calculated lethal electric field thresholds; mean ± SEM; One-way ANOVAs with Tukey’s post hoc (* p<0.05, ** p<0.01, *** p<0.001, **** p<0.0001); n = 6.

**Table 1.**
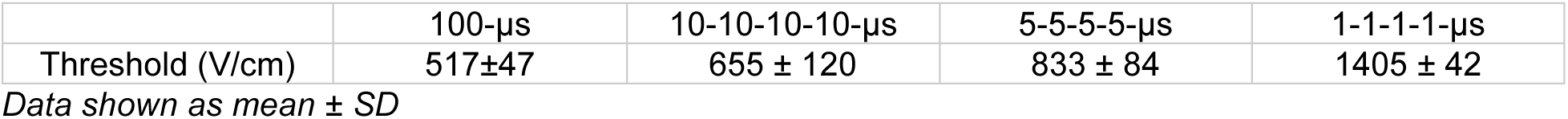
Cardiac Lethal Electric Field Thresholds.

### 2.6. Uniform electric fields applied using a parallel needle array validate bioimpedance saturation and lethal thresholds

Electroporaiton thresholds are an instrinsic property of the tissue that is waveform dependent, but is agnostic to the applied voltage, current, or electrode configurations^30^. To validate the lethal electric field thresholds, we utilized an 8-electrode parallel needle array with 1-cm spacing to deliver uniform electric fields for a subset of our characterized waveforms (1 and 10 µs). Voltages of 750, 1250, and 2500 V were delivered across the cardiac tissue to generate electric fields above and below our calculated lethal electric field thresholds. For 2500 V applied across 1 cm of spacing, we generated measurable leasions, while we could not find lesions for the sub-electroporative treatments (**Figure 6A**). The electrode geometry was then reconstucted within COMSOL as described above with 2500 V applied between the electrode arrays. A contour was defined using the calculated lethal electric field thresholds above, with 94.6% and 95.3% overlap for the 1-1-1-1-µs x 50 and 10-10-10-10-µs x 5 waveforms, respectively. The tissues were then processed and analyzed to visualize the lesion on H&E (**Figure 6B**); we did not observe coagulative necrosis associated with thermal damage but observed a slight decrease in deep purple within the lesion associated cell death (**Figure 6C**).

**Figure 6.**
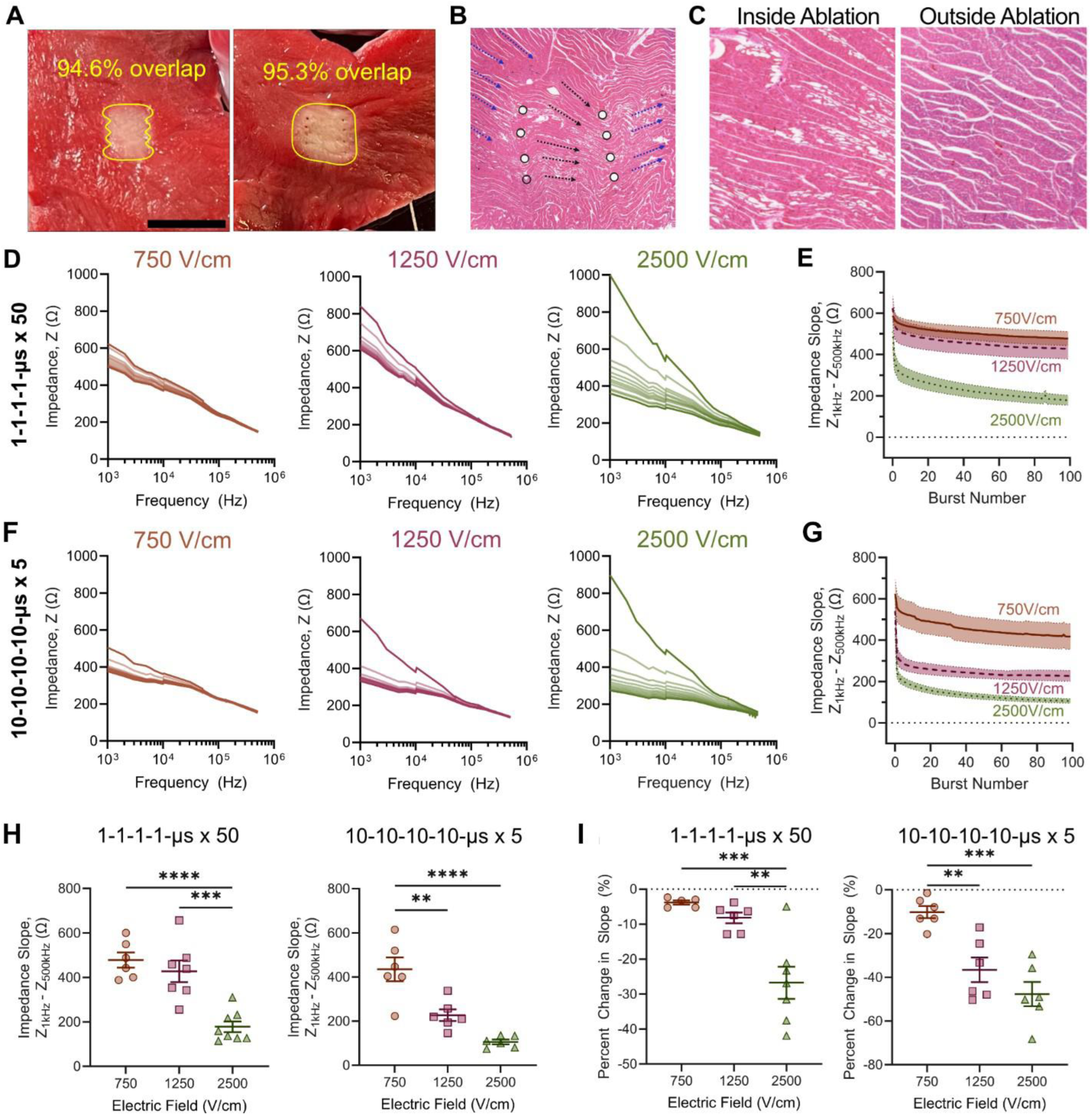
Ablation and bioimpedance assessment using parallel electrodes to deliver uniform electric fields. **A)** TCC staining of cardiac tissue slices ∼ 4 hours after pulsed field ablation with 2500 V across 1 cm of parallel needle electrodes. **B)** Hematoxylin & Eosin (H&E) demonstrates local fiber orientation and **C)** potential viability differences within and outside the lesion that is not attributed to thermal coagulative necrosis. Representative pre-treatment EIS, inter-burst EIS (every 10 bursts plotted), and post-treatment EIS measurements for the **D)** 1-µs and **F)** 10-µs waveforms over 100 bursts at 1 Hz. Difference in low-frequency and high-frequency impedance (slope) for pre-treatment (0), between applied bursts (1-99), and post-treatment (100) for the **E)** 1-µs and **G)** 10-µs waveforms; mean ± SEM; n = 6; data fit with an unrestricted 2-phase exponential decay. **H)** Impedance slops at burst 100 the 1-µs and 10-µs waveforms; mean ± SEM; n = 6; One-way ANOVAs with Tukey’s post hoc (*** *p <* 0.001, **** p < 0.0001). **I)** Percent change in impedance slope from pre-treatment to burst 1 (b_1_ – b_0_); mean ± SEM; One-way ANOVAs with Tukey’s post hoc (** *p <* 0.01, *** p < 0.001); n = 6.

For each treatment, FAST bioimpedance was also measured between each burst delivery. Similar to the 2-needle treatments, the slope initially dropped rapidly within the first packet, then more gradually with each subsequent packet (**Figure 6D-F**). The slope was also significantly lower for applied electric fields above the calculated lethal electric field threshold, when compared to applied electric fields below the calculated lethal electric field threshold (**Figure 6F**). Similarly, the percent change in slope was also greater for high electroporative treatments (**Figure 6G**). We attribute the small drop in bioimpedance for the sub-electroporative condition due to high electric field concentrations direclty adjacent to the small electrodes. These results help support that the measured change in impedance slope is due to the onset and progression of electroporation within the cardiac tissue, and validates the calculated lethal electric field threshold for our *ex vivo* experimental setup.

## 3. Discussion

Despite high acute pulmonary vein isolation rates, incomplete transmurally and lesion heterogeneity remain a concern, contributing to one-year recurrence rates of 20–45% for paroxysmal and persistent atrial fibrillation^38,39^. Since cell death following PFA occurs over hours, visualization of cardiac lesions and PV isolation is limited to expensive and invasive mapping systems. However, temporary cardiomyocyte stunning due to the sub-electroporative electric fields may confound maps directly after treatment^40^. Therefore, establishing alternative metrics for determining the formation of durable, homogeneous, and transmural ablations remains a critical barrier to long-term PFA success.

We demonstrate that cardiac tissue electrical impedance can be measured during PFA delivery and that tissue capacitance diminishes following a 2-phase decay during electroporation. Attempts to measure bioimpedance changes during treatments using MR Electrical Impedance Tomography have shown promise in a laboratory setting^41,42^; however, translating this approach to clinical application is limited due to the need for specialized equipment and MRI imaging during treatment. Other EIS approaches for measuring changes in impedance at the electrodes are similarly limited for clinical applications due to slow acquisition times (>1min)^43^. To address this, our group previously developed FAST to integrate with current pulsing systems to both monitor and inform treatment delivery in real-time^33,34^.

Our data suggest that residual capacitance remains within the tissue and would not be removed if we continued to deliver pulses. Since every sample generated contiguous lesions, we do not attribute the plateaued “slope” to intact non-electroporated cells within the lesion. Therefore, we propose defining electroporation saturation, the condition where further changes to the cell cannot be induced via electroporation, as the point at which the burst-to-burst percent change in impedance slope falls below a predefined threshold for successive bursts. This approach reflects the diminishing rate of capacitance change rather than assuming complete loss of capacitive behavior, aligning with the biophysical reality that the cell membrane does not disappear during permeabilization. This metric would then be waveform-agnostic, does not require curve fitting or post hoc modeling, and could be implemented for real-time analysis. Since many commercial systems have established treatment protocols, this metric could determine when saturation occurs by estimating cumulative effects. The cumulative effects can then potentially be correlated with catheter intracardiac electrogram measurements to provide additional data for deciding if cardiac tissue is reversibly or irreversibly electroporation, helping elucidate the extent of stunning beyond the lesion and timescale following pulse applications. Previous work in liver has suggested that bioimpedance measurements can be utilized to distinguish reversible and irreversible electroporation^44^, supporting this potential approach.

A key finding of this work is that electroporation-induced changes in tissue electrical properties effectively homogenize myocardial anisotropy during PFA delivery. While cardiac tissue is inherently partially anisotropic due to elongated cardiomyocytes and preferential fiber orientation, this anisotropy arises largely from intact cell membranes acting as capacitive barriers to current flow. Our inter-burst bioimpedance measurements demonstrate a rapid attenuation of membrane-associated capacitance during the initial bursts, consistent with widespread membrane permeabilization. Once pores are formed, current can traverse both intra- and extracellular domains, substantially reducing the directional dependence that governs low-field conduction. As a result, electric field distributions computed using homogenized conductivity models closely matched those generated using fully anisotropic tensors under electroporation conditions, and neither lesion morphology nor fitted lethal thresholds exhibited systematic dependence on fiber alignment. Our data and rationale support that pore formation reduces the tissue capacitance, which is the factor creating potential anisotropy. These findings provide experimental support for the hypothesis that dynamic electroporation effects dominate over baseline structural anisotropy during PFA, particularly in heterogeneous cardiac tissue where large regions of perfectly aligned fibers are uncommon. From a modeling perspective, this supports the use of homogenized conductivity formulations when electroporation-dependent conductivity is explicitly incorporated, reducing model complexity without sacrificing predictive accuracy.

Using this homogenized framework, we derived waveform-specific lethal electric field thresholds across pulse widths spanning 1 – 100 µs. The observed monotonic increase in lethal threshold with decreasing pulse width is consistent with electroporation theory, reflecting reduced membrane charging efficiency at shorter pulse durations and the need for higher field strengths to achieve irreversible permeabilization. Importantly, the thresholds reported here are higher than many values assumed in prior cardiac PFA modeling studies. By comprehensively characterizing electroporation-dependent conductivity in our previous work to accurately calculate the electric field distribution, we purposefully set the grounds for calculating the waveform-specific lethal thresholds from experimentally measured lesion areas. The thresholds presented here are thus grounded in both tissue response and realistic electric field distributions. This distinction is important as the conductivity ultimately determines the electric field distribution and subsequent lethal threshold calculation. Previous characterization papers fit conductivity values with minimal information, potentially biasing the calculated thresholds to lower values. Furthermore, we disclose the waveforms to facilitate the possibility of cross-validation and additional verification.

Several limitations of this study should be acknowledged. First, experiments were performed in fresh *ex vivo* ventricular tissue, which lacks perfusion and longer follow up evaluations. Although this isolates electroporation effects from confounding physiological variables, it may influence thermal dissipation and post-treatment lesion evolution. Temperature was not directly measured during PFA delivery; however, the absence of visual charring, lack of high-frequency impedance increases, preservation of tissue architecture on histology, and strong agreement between measured and simulated lesion boundaries collectively suggest that thermal injury did not play a dominant role under the conditions studied. Second, needle electrodes were used to ensure consistent tissue contact and well-defined electric field distributions, whereas most clinical PFA catheters interact with the myocardium through the blood pool and exhibit more complex geometries. The force that a catheter is applied to the endocardium has a direct positive correlation to lesion depth due to tissue compression, which can be non-uniform and difficult to remove from threshold calculations. To address these limitations and validate the derived thresholds independent of electrode configuration, we utilized two percutaneous needle configurations, including a parallel needle array for uniform electric field delivery. Lesion formation occurred only when applied fields exceeded the calculated lethal thresholds, and the resulting lesions exhibited >94% overlap with predicted contours. Concurrent bioimpedance measurements during these experiments reproduced the same saturation behavior observed with monopolar needles, providing independent confirmation that the impedance metrics and lethal thresholds reflect intrinsic tissue electroporation rather than geometry-specific artifacts. Together, these validation experiments strengthen the generalizability of the proposed framework while clarifying its current experimental boundaries.

We demonstrate that bioimpedance can be monitored in fresh *ex vivo* tissue, so future work should utilize FAST (or equivalent rapid EIS techniques) to characterize cardiac bioimpedance profile with commercial PFA catheters pressed against the endocardium, to determine if electroporation effects can be measured *via* EIS during treatment and correlated to lesion outcomes. While the inserted needle electrodes allow for circumferential contact with cardiac tissue, most commercial catheters do not puncture the epicardium. Therefore, significant amounts of current travels through the blood pool, both requiring some level of perfusion and introducing another variable geometric domain with different reactions to electroporation. Radiofrequency ablation (RFA) work has used similar change-in-bioimpedance measurements for indicating successful lesion formation which supports real use cases with PFA^45^. Unlike RFA which typically uses focal point catheters, PFA confers a wide range of electrode designs including multi-electrode bipolar catheters for single-shot pulmonary vein isolation. Multi-electrode catheters can potentially sample broadband bioimpedance between different electrode arrays for isolating electrode contact while selectively gathering information on electroporation from the electrode pairs in contact.

## 4. Conclusion

This study demonstrates that cardiac electroporation can be quantitatively monitored during PFA delivery through inter-burst bioimpedance measurements. The rapid reduction and subsequent stabilization of impedance slope across waveforms indicate early electroporation saturation, enabling the use of simple, burst-resolved metrics to assess treatment progression without post hoc modeling. Experimental lesion analysis combined with nonlinear anisotropic electroporation modeling demonstrated that electroporation effectively homogenizes tissue anisotropy, reducing the impact of fiber orientation on electric field distributions and lesion geometry. Using this framework, we establish validated, waveform-specific lethal electric field thresholds for cardiac tissue and confirm their predictive value under uniform field conditions. Together, these findings support the use of bioimpedance-assisted monitoring and homogenized modeling approaches to improve the consistency, interpretability, and reproducibility of pulsed field ablation therapies.

## 5. Methods

### Determining treatment voltage for the different PFA waveforms

The 3D, time-dependent multi-tissue model was developed within COMSOL^TM^ Multiphysics 6.4, with the material properties given in Supplemental Table 1 and detailed previously^30^. The electric field distribution and Joule heating within the tissues were modeled using the AC/DC and heat transfer models, respectively. Current conservation was simulated using a modified Laplace equation under a quasi-static approximation:

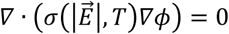

Where 𝜙 is the electric potential, and 𝜎 is the local electrical conductivity, which is dependent on the local electric field magnitude, 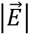, and temperature, T. Electroporation dependent conductivity changes were defined within the tissue using cardiac and waveform-specific conductivity curves:

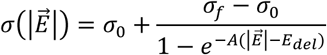

where 𝜎_0_ is the local bulk tissue conductivity, 𝜎_0_ is the local conductivity of electroporated tissue, and A and E_del_ are empirically determined sigmoidal fit parameters, with fit parameters recently derived for the high-frequency and conventional IRE waveforms used here^30^. Temperature was simulated using a modified Penne’s bioheat equation with the addition of Joule heating from the applied electric field:

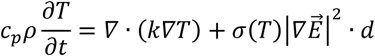

where 𝑐_𝑝_is the myocardium specific heat, is the tissue density, 𝑘 is the tissue thermal conductivity, and 𝑑 is the duty cycle for the applied burst. The local electrical conductivity of each tissue was dynamically updated by the increase in local temperature:

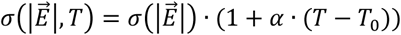

where 𝛼 is the thermal coefficient, which is the percent increase in electric conductivity from an increase in temperature, and 𝑇_0_ is the initial temperature, 37 °C. One electrode was defined as the voltage source, and the other was defined as the voltage sink. The voltage, V, was then parametrically swept within each computational model defining the waveforms. A cut plane was defined through the center of the electrode exposure, and the lesion area was then calculated by integrating the area within the IRE thresholds previously calculated *in vitro* ^30^. Computational model replicates for calculated areas used individual *in vitro* threshold replicate values.

### Delivery of Pulsed Field Ablation to Cardiac Tissue

Fresh porcine hearts were obtained from the University of Georgia Meat Science Technology Center (Athens, Georgia, USA). All experiments for a given heart were performed at the meat center within 20 minutes of death to maintain the viability of the tissue and body temperature (∼37-40 °C, measured) as described previously^30,32^. The heart was cut in half through the ventricle to directly expose the myocardium.

PFA was delivered through either two 18G diameter NanoKnife^TM^ (Angiodynamics) monopolar needles or a custom 8-electrode array with two parallel sets of four needles spaced 1 cm apart. The active needle exposure was 1-cm for both configurations and both were inserted ∼2 cm deep into the ventricle. The electrodes were inserted with the alignment being left and right, irrespective of position around the ventricle. Pulsed electric fields were generated using a custom high-voltage pulse generator (OmniPorator, VITAVE).

### Fourier Analysis Spectroscopy (FAST) Bioimpedance Measurements and Processing

Inter-burst electric impedance spectroscopy (EIS) was measured using the custom FAST board that controlled the timing and delivery of the FAST waveform and high-voltage pulsed electric fields^33,34^. Treatment was prefaced with an initial FAST waveform to sample to tissue before electroporation, followed by triggering the PFA burst. The board then delivered a FAST waveform 495 ms after every PFA burst for diagnostic measurement during delivery. The FAST waveform voltages and currents were monitored using a WaveSurfer 4024HD 5 GHz oscilloscope (Teledyne LeCroy) equipped with a 1000× attenuated high-voltage probe (DPB5700, Siglent) and a 10× attenuated current probe (3972, Pearson Electronics). A fixed sampling period of at least 80 ns was maintained to ensure that each 10ms-long FAST waveform was captured with 1.25 x 10^6^ points. A Fast Fourier Transform (FFT) then broke the concatenated recorded voltage and current waveforms into their frequency spectrums^33,46^. The data was then filtered to only keep points >10% the maximum voltage peak, commonly used to reduce noise from low power signals. The voltage at each remaining frequency was then divided by the current at the same frequency to get the impedance spectrum.

### Irreversible Electroporation Lesion Visualization and Measurements

The cardiac tissue was dwelled for 4 hours following PFA to allow for ablations to develop^32,47,48^ and then sectioned through the midline of the electrode (∼2.5 cm deep). The cardiac slices were then stained for viability using 2,3,5-Triphenyl tetrazolium chloride (TTC) in phosphate buffered saline (1.5% w/v; ThermoFisher, 10010023) for 10 minutes at room temperature with protection from direct light. We then imaged the lesions on a grid cutting mat using a digital camera. A custom holder was used to maintain 16.5 cm from the surface for every image. The images were then imported into ImageJ (National Institutes of Health) for analysis. The non-stained pale region identifying the IRE lesion was measured using the oval selection tool. The pixel value for the length of the 1 cm spacing was used to convert the measured pixel areas from ImageJ into centimeters squared.

### Anisotropic tissue simulations and homogenization fit calculations

Unlike previous modeling that scaled the entire conductivity curve by a constant factor for parallel and perpendicular directions^24^, the induction of electroporation and removal of membrane capacitance suggests an upper limit to the tissue conductivity^31^. We first calculated the conductivity with the fibers aligned with the electric field as:

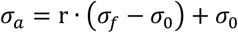

Where r is a chosen ratio from 0 to 1 to define the extent of anisotropy. The conductivity tensor for non-electroporated bulk conductivity, 𝜎_0_, was then calculated as:

|  |  |  |
| --- | --- | --- |
| $\sigma_{11} = \sigma_a \cdot \cos^2(\theta) + \sigma_0 \cdot \sin^2(\theta)$ | $\sigma_{12} = \cos(\theta) \sin(\theta) \cdot (\sigma_a - \sigma_0)$ | 0 |
| $\sigma_{21} = \sigma_{12}$ | $\sigma_{22} = \sigma_a \cdot \sin^2(\theta) + \sigma_0 \cdot \cos^2(\theta)$ | 0 |
| 0 | 0 | $\sigma_0$ |

Where 𝜃 is the dominant fiber angle in the counterclockwise direction on the xy-plane. Following the electrical conductivity curve of the tissue was defined to incorporate the bulk conductivity as an input to the equation:

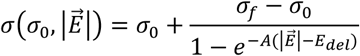

A symmetric tensor with the incorporation of the dynamic conductivity equation gives:

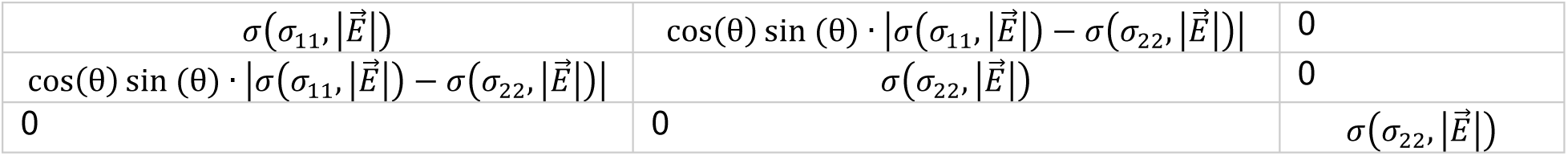

Note that both the bulk conductivity and the electroporated conductivity have defined skews. The conductivity was then adjusted within the time-dependent solution due to simulated heating as described above.

The homogenization fit was calculated by determining the electric field contours for each of the fit conditions (*i.e.,* isotropic, anisotropic, and isotropic with electroporation electroporation) with the same area measured for the lesion. The lesion image was imported into ImageJ, converted to 8-bit for auto-color-thresholding. The electric field contours were aligned so that the electrodes overlayed with those on the image. A freehand trace was drawn for the area that overlapped between the lesion and each contour. The pixel value for the overlapping area was then divided by the total pixel value for the lesion.

### Statistical Analyses

Statistical analyses were performed using GraphPad Prism 10.5.0 (GraphPad Software, Boston), with the specific tests used outlined within the figure legends. Post-experimental power analyses were performed using G*Power 3.1 (Heinrich Heine Universität, Düsseldorf)^49^.

## Data Availability Statements

The data that supports the findings of this study are available within the article and its supplementary material. Further requests can be made to the corresponding author.

## Declaration of competing interests

The authors declare the following financial interests/personal relationships which may be considered as potential competing interests: The authors have patents related to the paper. R.V.D. receives royalty income from technologies he has invented. S.D. is the Chief Technology Officer (CTO) of Contour Medical.

## Acknowledgements and Funding

This research was supported in part by the NSF/FDA SBIR award (#2129626) and by the New York State Division of Science, Technology and Innovation (NYSTAR) under Contract No. C200124 through Cornell University’s Center for Life Science Enterprise. We would like to acknowledge Harl Ryan Crowe and the University of Georgia Edgar L. Rhodes Center for Animal & Dairy Science for their help in procuring whole, fresh porcine hearts. We would also like to thank Ram Anand Vadlamani and Maura Casciola for early discussion on experimental design and direction.

## Supplement

**Supplemental Table 1:** Myocardial parameters not provided elsewhere.

| Parameter | Symbol | Value | Reference |
| --- | --- | --- | --- |
| Myocardium specific heat | $c_p$ | $3686 [J \cdot kg^{-1} \cdot K^{-1}]$ | 50 |
| Myocardium density | $\rho$ | $1081 [kg \cdot m^{-3}]$ | 50 |
| Myocardium thermal conductivity | $k$ | $0.56 [W \cdot m^{-1} \cdot K^{-1}]$ | 50 |
| Tissue thermal coefficient | $\alpha$ | $0.015 [1 \cdot K^{-1}]$ | 51 |

## References

1. Jacobs, E., Rubinsky, B. & Davalos, R. Pulsed Field Ablation in Medicine: Irreversible Electroporation and Electropermeabilization Theory and Applications. Radiol. Oncol. (2025) doi:10.2478/raon-2025-0011.

2. Davalos, R. V, Mir, L. M. & Rubinsky, B. Tissue Ablation with Irreversible Electroporation. Ann. Biomed. Eng. 33, 223–231 (2005).

3. Arena, C. B., Sano, M. B., Rossmeisl, J. H., Caldwell, J. L., Garcia, P. A., Rylander, M. N. & Davalos, R. V. High-frequency irreversible electroporation (H-FIRE) for non-thermal ablation without muscle contraction. Biomed. Eng. Online 10, 102 (2011).

4. DeBruin, K. A. & Krassowska, W. Modeling electroporation in a single cell. I. Effects of field strength and rest potential. Biophys. J. 77, 1213–1224 (1999).

5. Neumann, E. & Rosenheck, K. Permeability changes induced by electric impulses in vesicular membranes. J. Membr. Biol. 10, 279–290 (1972).

6. Edd, J. F., Horowitz, L., Davalos, R. V., Mir, L. M. & Rubinsky, B. In vivo results of a new focal tissue ablation technique: Irreversible electroporation. IEEE Trans. Biomed. Eng. 53, 1409–1415 (2006).

7. Al-Sakere, B., André, F., Bernat, C., Connault, E., Opolon, P., Davalos, R. V., Rubinsky, B. & Mir, L. M. Tumor Ablation with Irreversible Electroporation. PLoS One 2, (2007).

8. Koruth, J. S., Kuroki, K., Kawamura, I., et al. Pulsed Field Ablation Versus Radiofrequency Ablation: Esophageal Injury in a Novel Porcine Model. Circ. Arrhythm. Electrophysiol. 13, E008303 (2020).

9. Ekanem, E., Neuzil, P., Reichlin, T., et al. Safety of pulsed field ablation in more than 17,000 patients with atrial fibrillation in the MANIFEST-17K study. Nat. Med. 30, 2020–2029 (2024).

10. Reddy, V. Y., Gerstenfeld, E. P., Natale, A., et al. Pulsed Field or Conventional Thermal Ablation for Paroxysmal Atrial Fibrillation. New England Journal of Medicine 389, 1660–1671 (2023).

11. Ekanem, E., Reddy, V. Y., Schmidt, B., et al. Multi-national survey on the methods, efficacy, and safety on the post-approval clinical use of pulsed field ablation (MANIFEST-PF). Europace 24, 1256–1266 (2022).

12. Reddy, V. Y., Calkins, H., Mansour, M., et al. Pulsed Field Ablation to Treat Paroxysmal Atrial Fibrillation: Safety and Effectiveness in the ADMIRE Pivotal Trial. Circulation (2024) doi:10.1161/CIRCULATIONAHA.124.070333.

13. Reddy, V. Y., Mansour, M., Calkins, H., et al. Pulsed Field vs Conventional Thermal Ablation for Paroxysmal Atrial Fibrillation: Recurrent Atrial Arrhythmia Burden. J. Am. Coll. Cardiol. 84, 61–74 (2024).

14. Verma, A., Haines, D. E., Boersma, L. V., et al. Pulsed Field Ablation for the Treatment of Atrial Fibrillation: PULSED AF Pivotal Trial. Circulation 147, 1422–1432 (2023).

15. Anter, E., Mansour, M., Nair, D. G., et al. Dual-energy lattice-tip ablation system for persistent atrial fibrillation: a randomized trial. Nat. Med. 30, 2303–2310 (2024).

16. Duytschaever, M., De Potter, T., Grimaldi, M., et al. Paroxysmal Atrial Fibrillation Ablation Using a Novel Variable-Loop Biphasic Pulsed Field Ablation Catheter Integrated With a 3-Dimensional Mapping System: 1-Year Outcomes of the Multicenter inspIRE Study. Circ. Arrhythm. Electrophysiol. 16, E011780 (2023).

17. García-Bolao, I., Reddy, V. Y., Su, W. W., et al. Visualization of PFA During PVI With the Second-Generation Pentaspline Catheter. JACC Clin. Electrophysiol. (2025) doi:10.1016/j.jacep.2025.11.016.

18. Julian Chun, K. R., Miklavčič, D., Vlachos, K., Bordignon, S., Scherr, D., Jais, P. & Schmidt, B. State-of-the-art pulsed field ablation for cardiac arrhythmias: ongoing evolution and future perspective. Europace 26, (2024).

19. Koop, B. Fundamentals of System Design for Cardiac Pulsed Field Ablation: Optimization of Safety, Efficacy, and Usability. PACE - Pacing and Clinical Electrophysiology Preprint at 10.1111/pace.15120 (2025).

20. Garrott, K., Bifulco, S., Ramirez, D. & Koop, B. Lesion Formation in Cardiac Pulsed-Field Ablation: Acute to Chronic Cellular Level Changes. PACE - Pacing and Clinical Electrophysiology Preprint at 10.1111/pace.15154 (2025).

21. Yamashita, K., Kikuchi, Y., Yoshiyama, K., Daiki, |, Yosuke, K. |, | M. & Onodera, K. Novel Impedance-Guided Contact Mapping Technique for the Circular Multielectrode Pulsed-Field Ablation Catheter. Clin. Case Rep. (2026) doi:10.1002/ccr3.72142.

22. Fortuna, M., Combes, N., Combes, S., Cardin, C., Voglimacci-Stephanopoli, Q., Boveda, S. & Albenque, J. P. Posterior mitral isthmus ablation using a balloon-in-basket pulsed-field multielectrode catheter. HeartRhythm Case Rep. (2026) doi:10.1016/j.hrcr.2026.02.015.

23. Hirata, S., Nagashima, K., Watanabe, R., et al. Exploratory Assessment of Field-Tag–Based Catheter–Tissue Contact and Acute Lesion Surrogates During Pulsed Field Ablation. Heart Rhythm *O2* (2026) doi:10.1016/j.hroo.2026.02.003.

24. Kos, B., Mattison, L., Ramirez, D., Cindrič, H., Sigg, D. C., Iaizzo, P. A., Stewart, M. T. & Miklavčič, D. Determination of lethal electric field threshold for pulsed field ablation in ex vivo perfused porcine and human hearts. Front. Cardiovasc. Med. 10, (2023).

25. Kwon, H., Guasch, M., Nagy, J. A., Rutkove, S. B. & Sanchez, B. New electrical impedance methods for the in situ measurement of the complex permittivity of anisotropic skeletal muscle using multipolar needles. Sci. Rep. 9, (2019).

26. Sato, H., Nakamura, T., Kusuhara, T., Kenichi, K., Kuniyasu, K., Kawashima, T. & Hanayama, K. Effectiveness of impedance parameters for muscle quality evaluation in healthy men. Journal of Physiological Sciences 70, (2020).

27. Šmerc, R., Miklavčič, D. & Mahnič-Kalamiza, S. Skeletal muscle death from the perspective of electrical impedance as evidenced by experiment and numerical modelling. Comput. Biol. Med. 197, (2025).

28. Čorović, S., Županič, A., Kranjc, S., Al Sakere, B., Leroy-Willig, A., Mir, L. M. & Miklavčič, D. The influence of skeletal muscle anisotropy on electroporation: in vivo study and numerical modeling. Med. Biol. Eng. Comput. 48, 637–648 (2010).

29. Wong, J. & Kuhl, E. Generating fibre orientation maps in human heart models using Poisson interpolation. Comput. Methods Biomech. Biomed. Engin. 17, 1217–1226 (2014).

30. Jacobs, E., Santos, P. & Davalos, R. Comprehensive characterization of waveform-dependent cardiac tissue electroporation for pulsed field ablation. Biosens. Bioelectron. 289, (2025).

31. Castellvi, Q. & Ivorra, A. Computational Multiscale Modeling of Pulsed Field Ablation Considering Conductivity and Damage Anisotropy Reveals Deep Lesion Morphologies. Int. J. Numer. Method. Biomed. Eng. 41, (2025).

32. Jacobs, E., Santos, P. & Davalos, R. Effects of Interphase and Interpulse Delays on Tissue Impedance and Pulsed Field Ablation. Ann. Biomed. Eng. (2025) doi:10.1007/s10439-025-03757-4.

33. Lorenzo, M. F., Bhonsle, S. P., Arena, C. B. & Davalos, R. V. Rapid Impedance Spectroscopy for Monitoring Tissue Impedance, Temperature, and Treatment Outcome During Electroporation-Based Therapies. IEEE Trans. Biomed. Eng. 68, (2021).

34. Campelo, S. N., Jacobs, E. J., Aycock, K. N. & Davalos, R. V. Real-Time Temperature Rise Estimation during Irreversible Electroporation Treatment through State-Space Modeling. Bioengineering 2022, Vol. 9, Page 499 9, 499 (2022).

35. Ivorra, A. Bioimpedance Monitoring for Physicians: An Overview. (2002).

36. Abasi, S., Aggas, J. R., Garayar-Leyva, G. G., Walther, B. K. & Guiseppi-Elie, A. Bioelectrical Impedance Spectroscopy for Monitoring Mammalian Cells and Tissues under Different Frequency Domains: A Review. ACS Measurement Science Au vol. 2 495–516 Preprint at 10.1021/acsmeasuresciau.2c00033 (2022).

37. Jacobs, E., Campelo, S. N., Charlton, A., Altreuter, S. & Davalos, R. V. Characterizing reversible, irreversible, and calcium electroporation to generate a burst-dependent dynamic conductivity curve. Bioelectrochemistry 155, 108580 (2024).

38. Queiroga, F., Rivera, A., Consoli, L. N., et al. Pulsed field versus thermal ablation for atrial fibrillation: A Bayesian meta-analysis. Heart Rhythm (2026) doi:10.1016/j.hrthm.2026.02.028.

39. Bolhuis, R. E., Bax, I. N., van Dijk, V. F., Balt, J. C., Wijffels, M. C. E. F., Boersma, L. V. A. & Liebregts, M. Pulsed Field Ablation for Atrial Fibrillation with a Balloon-in-Basket System under Conscious Sedation: A Retrospective Feasibility and Safety Analysis. Heart Rhythm (2026) doi:10.1016/j.hrthm.2026.02.011.

40. Cespón-Fernández, M., Nakasone, K., Pannone, L., et al. First Clinical Experience With Reversible Electroporation Mapping in Atrial Flutter. Circ. Arrhythm. Electrophysiol. (2026) doi:10.1161/CIRCEP.125.014359.

41. Granot, Y., Ivorra, A., Maor, E. & Rubinsky, B. In vivo imaging of irreversible electroporation by means of electrical impedance tomography. Phys. Med. Biol. 54, 4927–4943 (2009).

42. Kranjc, M., Kranjc, S., Bajd, F., Serša, G., Serša, I. & Miklavčič, D. Predicting irreversible electroporation-induced tissue damage by means of magnetic resonance electrical impedance tomography. Sci. Rep. 7, (2017).

43. Wang, C., Zhang, Y., Lin, F., et al. Bioimpedance assessment method based on back propagation neural network for irreversible electroporation of liver tissue. Sci. Rep. 15, (2025).

44. Ivorra, A. & Rubinsky, B. In vivo electrical impedance measurements during and after electroporation of rat liver. Bioelectrochemistry 70, 287–295 (2007).

45. Qian, P. C., Nguyen, D. M., Barry, M. A., Tran, V., Lu, J., Thiagalingam, A., Thomas, S. P. & McEwan, A. Optimizing Impedance Change Measurement During Radiofrequency Ablation Enables More Accurate Characterization of Lesion Formation. JACC Clin. Electrophysiol. 7, 471–481 (2021).

46. Bhonsle, S., Lorenzo, M. F., Safaai-Jazi, A. & Davalos, R. V. Characterization of nonlinearity and dispersion in tissue impedance during high-frequency electroporation. IEEE Trans. Biomed. Eng. 65, 2190–2201 (2018).

47. Younis, A., Demian, J., Bifulco, S., et al. Factors Influencing Lesion Titration in a Monopolar Pulsed-Field Ablation Point Catheter: A Preclinical Study. Circ. Arrhythm. Electrophysiol. (2025) doi:10.1161/CIRCEP.125.014157.

48. Nakagawa, H., Farshchi-Heydari, S., Sugawara, M., et al. Real-Time Prediction of Irreversible Lesion Size During Pulsed Field Ablation: Prospective Validation of a Novel Ablation Index Based on Contact Force and Number of Applications in a Swine Beating Heart Model. Circ. Arrhythm. Electrophysiol. 18, (2025).

49. Faul, F., Erdfelder, E., Lang, A.-G. & Buchner, A. G*Power 3: A flexible statistical power analysis program for the social, behavioral, and biomedical sciences. Behavior Reserach Methods 39, 175–191 (2007).

50. Gabriel, S., Lau, R. & Gabriel, C. The dielectric properties of biological tissues: Measurements in the frequency range 10 Hz to 20GHz.

51. Jacobs, E., Campelo, S., Aycock, K., Yao, D. & Davalos, R. V. Spatiotemporal estimations of temperature rise during electroporation treatments using a deep neural network. Comput. Biol. Med. 107019 (2023) doi:10.1016/j.compbiomed.2023.107019.

